# Selective inhibition of somatostatin-positive dentate hilar interneurons induces age-related cellular changes and cognitive dysfunction

**DOI:** 10.1101/2022.10.05.511002

**Authors:** Jinrui Lyu, Rajasekar Nagarajan, Maltesh Kambali, Muxiao Wang, Uwe Rudolph

## Abstract

The cellular basis of age-related impairments of hippocampal function is not fully understood. In order to evaluate the role of somatostatin-positive (Sst^+^) interneurons in the dentate gyrus hilus in this process, we chemogenetically inhibited Sst^+^ interneurons in the dentate gyrus (DG) hilus. Chronic chemogenetic inhibition (CCI) of these neurons resulted in increased c-Fos staining in the DG hilus, a decrease in the percentage of Gad67- and of Sst-expressing interneurons in the DG, and increased microglial activation in DG, CA3, and CA1. Total dendritic length and spine density were reduced in DG and CA1, suggesting reduced dendritic complexity. Behaviorally, the recognition index in an object recognition task and the percentage of spontaneous alternations in the Y maze were decreased, while in both initial and reversal learning in the Morris water maze the latencies to find the hidden platform were increased, suggesting cognitive dysfunction. Our findings establish a causal role for a reduced function of Sst^+^ interneurons in the DG hilus for cognitive decline and suggest that this reduced function may contribute to age-related impairments of learning and memory. Furthermore, our CCI mice may represent a cellularly defined model of hippocampal aging.

**SIGNIFICANCE STATEMENT:** Neuronal circuits and cellular processes underlying age-related cognitive dysfunction are not well understood. We observed that chronic chemogenetic inhibition of a defined cell type, somatostatin-positive interneurons in the dentate gyrus hilus, which have previously been found to be associated with cognitive dysfunction in aged rodents, is necessary and sufficient to elicit changes in expression of interneuronal markers, an increase in the activity of dentate gyrus granule cells, increased microglial activation across the entire hippocampus and an impairment of learning and memory-related tasks. Thus, inhibition of somatostatin-positive interneurons in the dentate gyrus hilus replicates changes that are also seen with normal aging, representing a novel cellularly defined animal model of hippocampal aging.

## INTRODUCTION

Aging is an inevitable, complex, and multifactorial process. Cognitive decline, such as memory loss, is one of the most prevalent concerns in the aging population. The hippocampus plays a vital role in cognition, learning, and memory in the mammalian brain ^1–3^. Importantly, the number of GABAergic interneurons in the hippocampus is reduced by normal aging ^4–6^, and hilar interneuron vulnerability has been shown to be correlated with age-regulated memory impairment ^6^. Optogenetic inhibition of hilar GABAergic interneuron activity impairs spatial memory ^7^. Moreover, many studies showed that aging in rodents is associated with DG and CA3 hyperactivity ^8,9^, CA1 hypoactivity ^9–11^, and reduced glutamatergic and GABAergic signaling in the hippocampus ^12^.

The dentate gyrus (DG), the first input-receiving region in the trisynaptic hippocampal pathway, is located between the entorhinal cortex and the CA3 region of the hippocampus. It receives information from the entorhinal cortex, projects to the CA3 pyramidal neurons, which then project to CA1 pyramidal neurons. DG granule cells encode spatial and contextual information ^13^. The tonic inhibition of DG granule cells, which are main regulators of hippocampal neuronal activity, controls pattern separation by distinguishing overlapping interferences into distinct and nonoverlapping information ^14^. α5 subunit-containing GABA_A_ receptors (α5-GABA_A_R), which are strongly expressed in the hippocampus, where they are primarily located extrasynaptically, mediate tonic inhibition in DG granule cells, CA1 and CA3 pyramidal cells ^14,15^. Mice with a conditional knockout of α5-GABA_A_R in dentate gyrus granule cells, which display a reduced tonic inhibition but preserved phasic inhibition, perform worse in many behavioral tasks associated with high memory interference. These mice displayed increased c-Fos staining in DG and CA3, consistent with hyperactivity ^14^. Interestingly, hyperactivity of DG and CA3 have also been reported to be linked to age-related memory decline in aging humans ^16^.

Somatostatin (Sst)-expressing interneurons are a subpopulation of GABAergic interneurons. Sst^+^ interneurons regulate hippocampal networks through dendritic inhibition of the DG circuitry ^17^. In the ApoE4 KI model of Alzheimer’s disease (AD) that showed learning and memory deficits, the numbers of hilar somatostatin-positive interneurons were decreased age-dependently ^18^. After receiving compounds that reduced hippocampal activity, both aged rats and human patients with amnestic mild cognitive impairment improved their memory ^19,20^. Moreover, age-related cognitive deficits have been found to be correlated with a decrease in somatostatin-positive interneurons in the dentate gyrus hilus in rats ^6^, however, a causal relationship has not been demonstrated.

In this study, we examined whether suppression of the activity of hilar somatostatin-positive (Sst^+^) interneurons is sufficient to induce defined learning and memory deficits, and whether this would occur only with chronic or also with acute inhibition. Our primary hypothesis was that only chronic inhibition of Sst^+^ interneurons results in learning and memory deficits. We tested this hypothesis by generating and analyzing mice in which these neurons could be chemogenetically silenced by clozapine. If reduction of activity of Sst^+^ interneurons in the dentate hilus resulted in cognitive dysfunction, the mice with such changes would potentially also represent a novel mouse model of age-related cognitive decline.

## RESULTS

The goal of this study was to determine whether a factor that has been observed in aged animals, i.e., a decline in the number of somatostatin-positive interneurons in the dentate gyrus hilus, is sufficient to induce cognitive decline and may thus underlie age-related cognitive dysfunction. Aging being a chronic process, we compared the effects of chronic and acute chemogenetic inhibition (CCI and ACI, respectively) of these neurons at the cellular and behavioral levels in order to determine whether chronic inhibition is required for these changes (**Fig. 1**).

**Figure 1.**
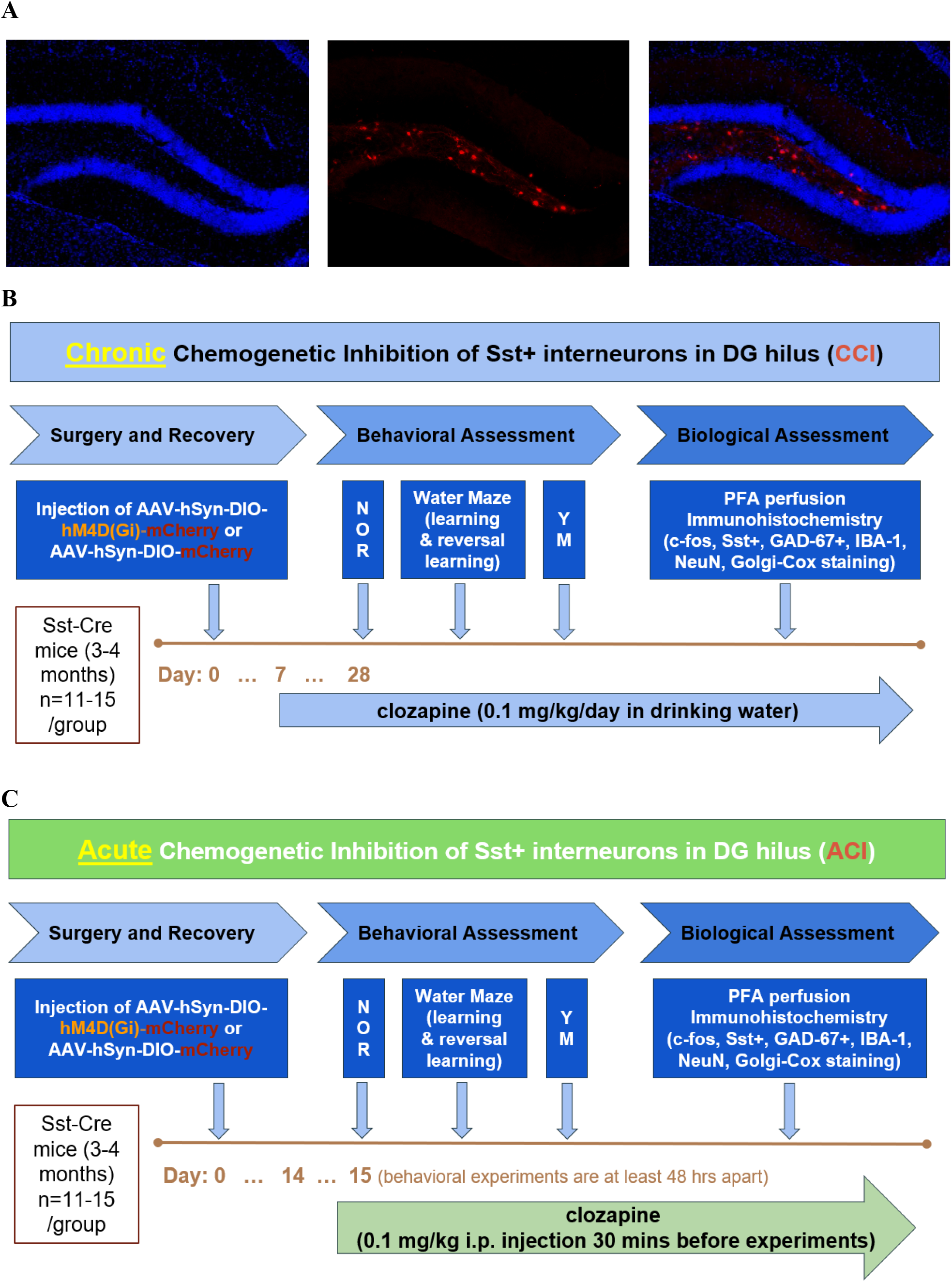
Experimental design of DG hilus somatostatin-positive (Sst^+^) cell manipulation. ***A***, Immunofluorescence staining of coronal sections from an AAV-mCherry mouse showing DAPI counterstain (left: DAPI; middle: mCherry expression; right: merged image).” ***B***, Top, Chronic chemogenetic inhibition of DG hilus Sst^+^ cells. Bottom, Acute chemogenetic inhibition of DG hilus Sst^+^ cells.

### Cellular activity, interneuronal markers and microglial activation

In order to assess how a reduced activity of Sst^+^ interneurons affects the activity of principal neurons in hippocampal subregions, we performed c-Fos staining in DG hilus, DG granule cell layer (GCL), CA3, and CA1 subregions of the hippocampus.

We observed that in mice with a chronic chemogenetic inhibition of Sst^+^ interneurons in the dentate gyrus (CCI) c-Fos staining in total DG (Diff=13.65, t=5.977) and in DG hilus (Diff=14.22, t=6.225) was increased (p<0.001, F_(4,50)_ =224.43, two-way ANOVA), consistent with hyperactivity, however, c-Fos staining in CA3 and CA1 was unaltered (p>0.05, **Fig. 2A**, top panel). In mice with an acute chemogenetic inhibition of Sst^+^ in the dentate gyrus (ACI), c-Fos staining also increased in DG hilus (Diff=8.273, t=3.096) and DG GCL (Diff=7.584, t=2.838) (p < 0.05, F_(4,50)_ =133.77, two-way ANOVA), but not in CA3 and CA1 (p>0.05, **Fig. 2A**, bottom panel). This suggests that both chronic and acute DG hilus-selective inactivation of Sst^+^ interneurons leads to increased c-Fos^+^ expression in the DG, possibly due to reduced tonic inhibition due to lack of an inhibitory input. CCI manipulation resulted in a 50% increase of c-Fos staining in the DG region (p<0.001, F_(4,60)_=38.35, Diff= 149.6, t=14.50, two-way ANOVA) with a 51% increase in the DG hilus (p<0.001, Diff=50.82, t=4.928) (**Fig. 2B**, left panel) while ACI manipulation resulted a 29% increase in the DG region (p<0.001, F_(4,60)_ =5.967, Diff=28.87, t=4.779) with a 34% increase in the DG hilus (p<0.001, Diff=34.21, t=5.664), and a 24% increase in the DG GCL (p<0.001, Diff=24.66, t=4.083), which is an indication of hyperactivity in the DG region (**Fig. 2B**, right panel).

**Figure 2.**
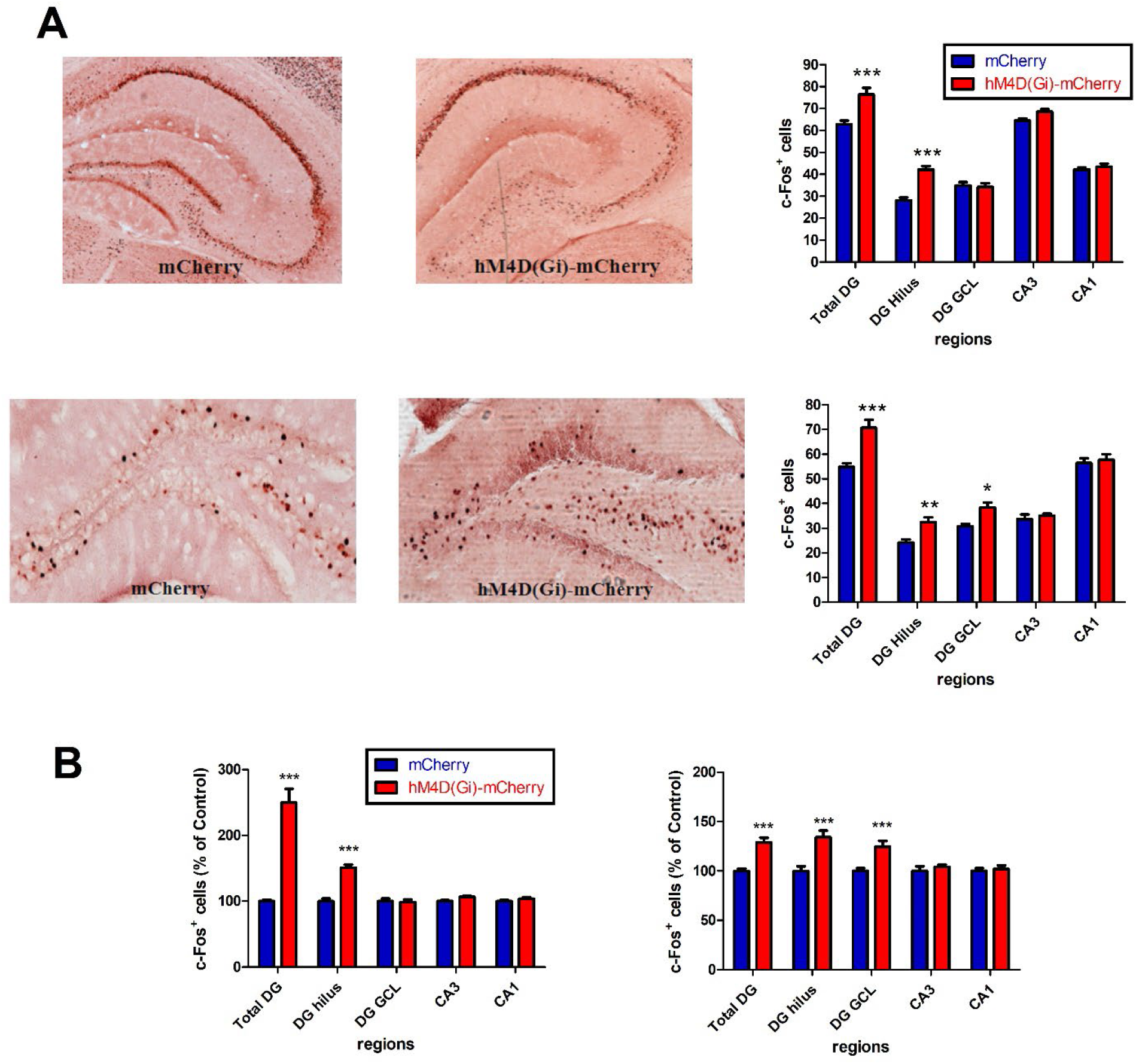
c-Fos^+^ cell counts in hippocampal subfields (Total DG, DG hilus, DG granule cell layer (DG GCL), CA3, CA1). ***A***, Top, Representative sections showing c-Fos^+^ expression in AAV-mCherry mice and AAV-hM4D(Gi)-mCherry mice during chronic chemogenetic inhibition (CCI). Bottom, Representative sections showing c-Fos^+^ expression in AAV-mCherry and AAV-hM4D(Gi)-mCherry mice during acute chemogenetic inhibition (ACI). ***B***, Left, Estimated density of c-Fos^+^ cells in AAV-hM4D(Gi)-mCherry mice expressed as percentage of AAV-mCherry mice during CCI. Right, Estimated density of c-Fos^+^ cells in AAV-hM4D(Gi)-mCherry mice expressed as percentage of AAV-mCherry mice during ACI. *p<0.05, **p<0.01, ***p<0.001 compared with the corresponding AAV-mCherry group.

We then assessed the influence of chronic and acute chemogenetic inhibition of Sst^+^ interneurons on the interneuronal markers Sst and GAD-67. While CCI mice displayed a significant decrease of Sst staining in DG (p<0.001, F_(4,50)_ =114.1, Diff=-10.53, t=7.43; hilus: p<0.001, Diff=-5.694,t=4.023; GCL: p<0.001, Diff=-4.833,t=3.415) (**Fig. 3A**, top panel), GAD-67^+^ was significantly decreased only in DG hilus (p<0.01, F_(4,50)_ =70.36, Diff= -9.540, t=3.675) (**Fig. 3B**, top panel). In ACI mice, no significant change was observed for both Sst^+^ and GAD-67^+^ staining (p > 0.05, two-way ANOVA) (**Fig. 3A, B**, bottom panels). We observed a loss of 51% of Sst^+^ interneurons in the DG region (p<0.001, F_(4,60)_=16.72, Diff=-51.26, t=6.985, two-way ANOVA) with a 75% reduction in the DG hilus (p<0.001, Diff=-74.65, t=10.17) and a 37% reduction in the DG GCL p<0.001, Diff=-37.44, t=5.102) with the CCI treatment (**Fig. 3C**, left panel). For GAD-67^+^ neurons, there is a 68% loss in the DG (p<0.001, F_(4,60)_=9.717, Diff=-67.89, t=7.069, two-way ANOVA) with a 64% loss in the DG hilus (p<0.001, Diff=-64.14, t=6.679; 74% loss in the DG GCL p<0.001, Diff=-74.07, t=7.713) (**Fig. 3C**, right panel).

**Figure 3.**
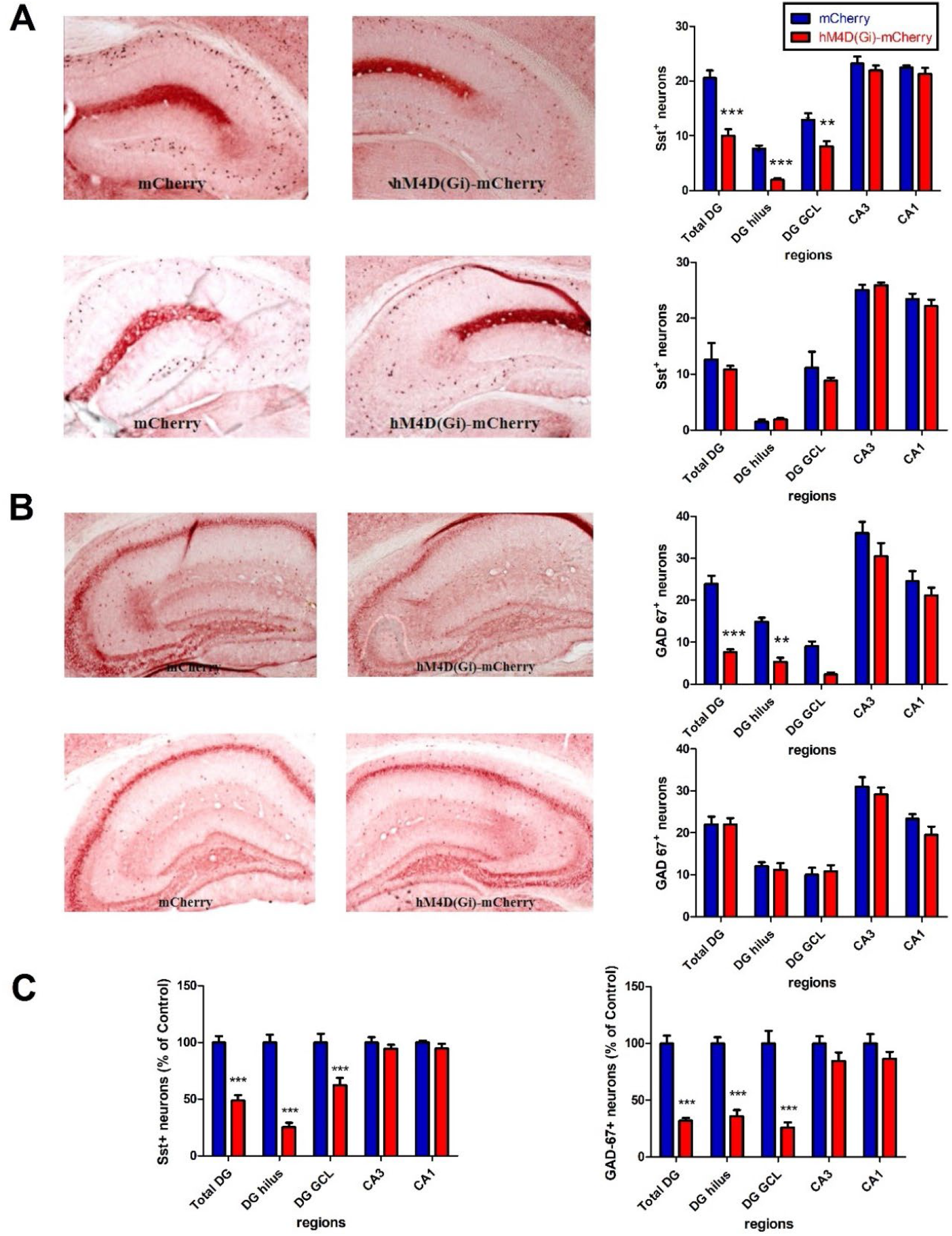
Sst^+^ neuron and GAD67^+^ neuron counts in hippocampal subfields. ***A***, Top, Representative sections showing Sst expression in AAV-mCherry and AAV-hM4D(Gi)-mCherry mice after CCI. Bottom, Sst expression during ACI. ***B***, Top, Representative sections showing GAD-67 expression in AAV-mCherry and AAV-hM4D(Gi)-mCherry mice during CCI. Bottom, GAD-67 expression during ACI. ***C***, Left, Estimated density of Sst^+^ neurons in AAV-hM4D(Gi)-mCherry mice expressed as percentage of AAV-mCherry mice during CCI. Right, Estimated density of GAD-67^+^ cells in AAV-hM4D(Gi)-mCherry mice expressed as percentage of AAV-mCherry mice during CCI. *p<0.05, **p<0.01, ***p<0.001

We used IBA1 (<u>I</u>onized Ca^2+^-<u>b</u>inding <u>a</u>dapter protein 1) as a cellular marker for microglial activity to investigate whether chemogenetic inhibition led to hippocampal microglial activation by determining average cell body size in CCI and ACI mice. Whereas ACI did not result in changes in the microglial cell body size (**Fig. 4B**, right panel, p> 0.05), CCI resulted in an increased average microglial cell body size in DG (q=5.821, p<0.001, one-way ANOVA), CA3 (q=4.757, p<0.01), and CA1 (q=4.480, p<0.01) (**Fig. 4B**, left panel). As a control experiment, we stained for NeuN^+^, which is expressed by all neurons. Neither ACI (F_(4,25)_=0.8259, p>0.05, two-way ANOVA) nor CCI (F_(4,25)_=0.1460 p>0.05, two-way ANOVA) resulted in a change in the number of neurons (**Fig. 4A**). These data suggest a reduced expression of the markers Sst^+^ and GAD-67^+^ in DG hilar interneurons and in the DG granule cell layer in the absence of a loss of neurons, and that this change results in increased hippocampal microglial activation in DG, CA3 and CA1. As it has been reported recently that microglia is involved in negative feedback control of neuronal activity ^28^, this microglial activation might potentially represent a compensatory mechanism and would then be consistent with insufficient tonic inhibition via GABA_A_Rs.

**Figure 4.**
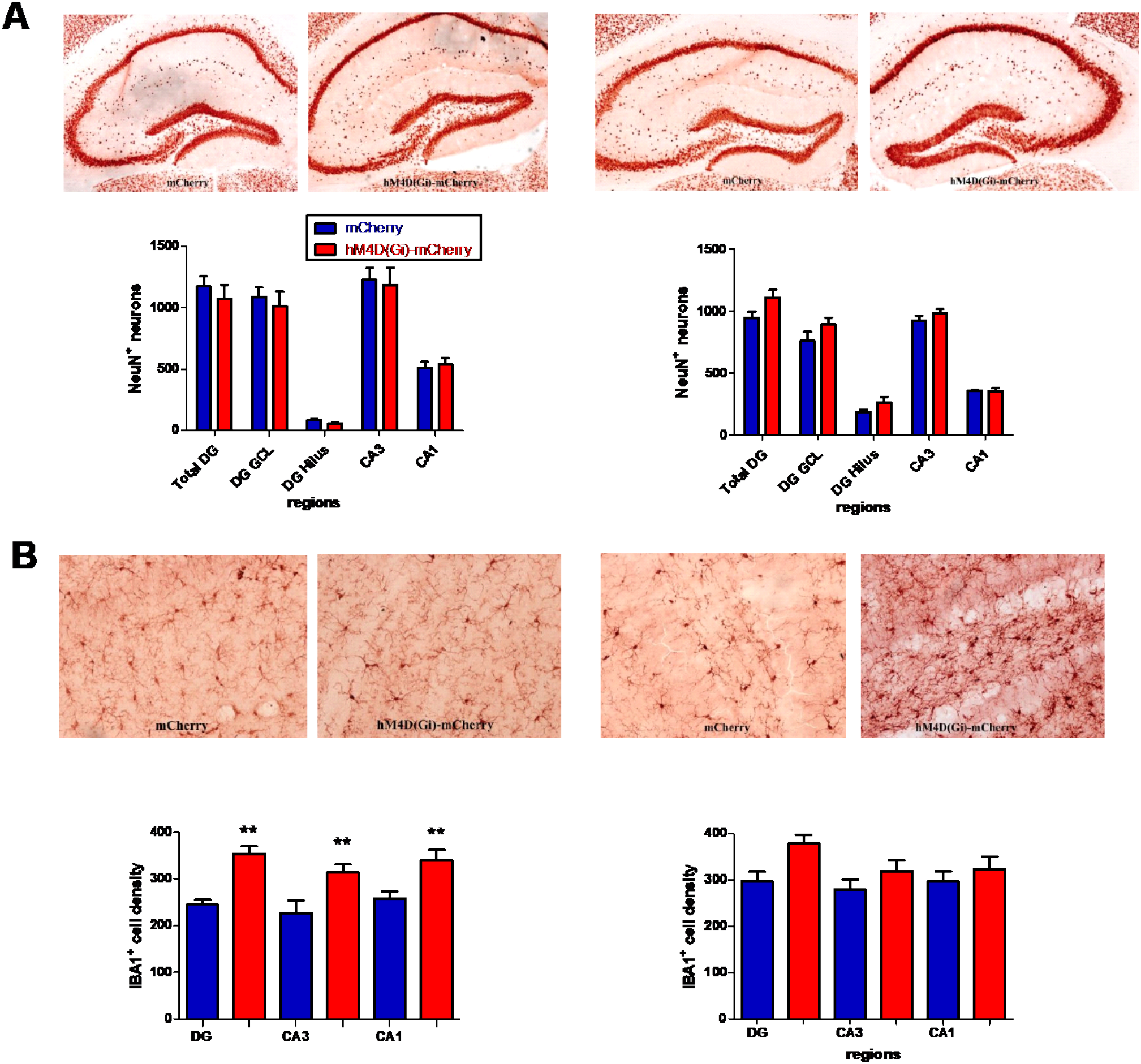
NeuN neuron counts and IBA1 cell density analysis in hippocampal subfields. ***A***, Left, Representative sections showing NeuN (total neurons) expression in mCherry and hM4D(Gi)-mCherry mice during CCI. Right, NeuN^+^ neurons expression during ACI. ***B***, Left, Representative sections showing IBA1^+^ cell density in mCherry and hM4D(Gi)-mCherry mice during CCI. Right, Representative sections showing IBA1^+^ cell density in mCherry and hM4D(Gi)-mCherry mice during ACI. *p<0.05, **p<0.01, ***p<0.001

### Spine density and dendritic length

To evaluate the impact of chronic and acute chemogenic inhibition of DG hilar Sst^+^ interneurons on hippocampal spine densities and spine length, we analyzed dendrites of granule cells located in the dentate gyrus and apical dendrites of CA1 pyramidal neurons (**Fig. 5A, B**). Four mouse brains were collected for Golgi-Cox staining from each treatment group. Quantification of dendritic length and spine density in DG and CA1 region was performed on 5-6 neurons per mouse using *Reconstruct* software.

**Figure 5.**
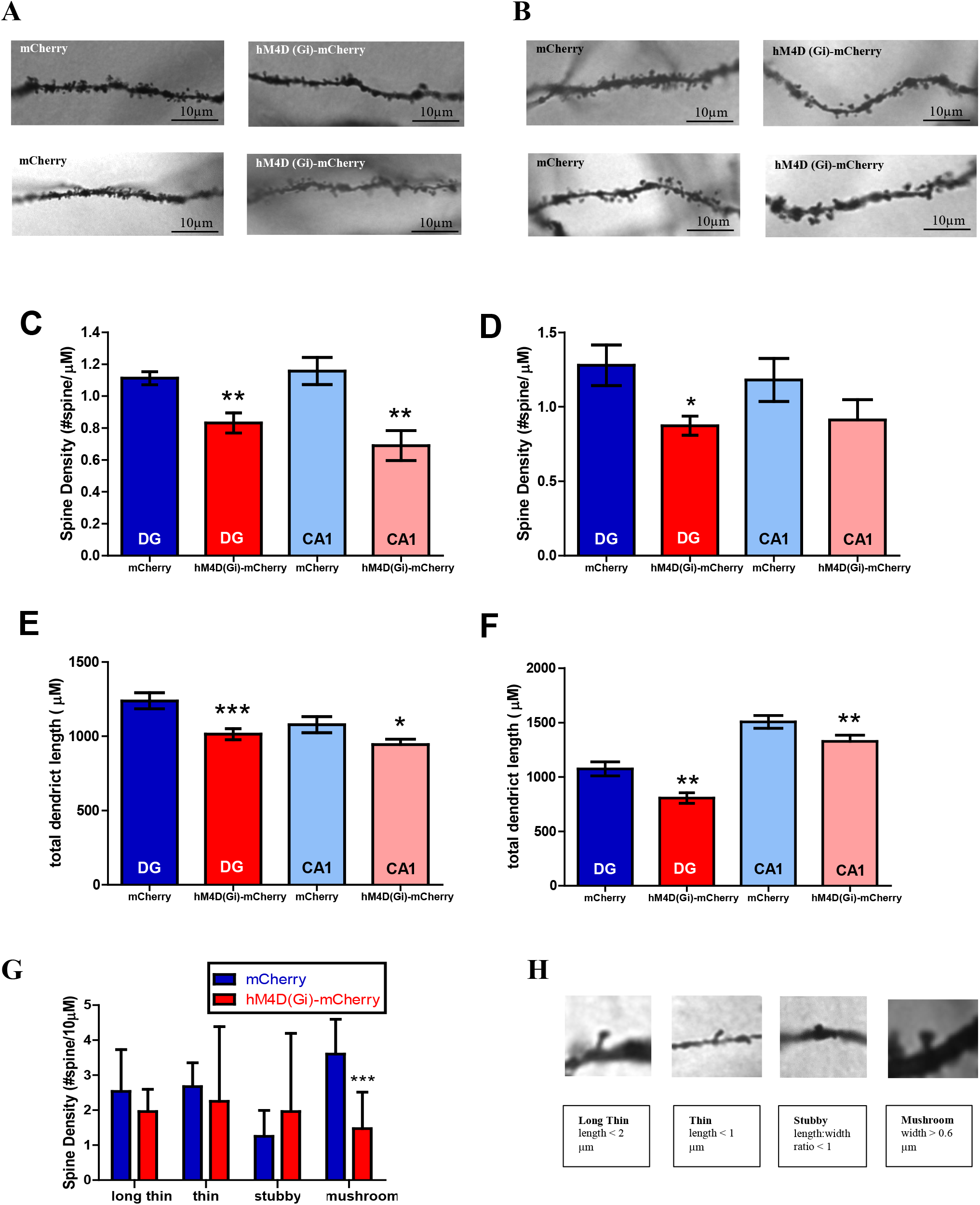
Morphological features of DG and CA1 regions in the hippocampus after CCI and ACI. ***A***, Top, Representative pictures of CCI in DG dendrite and spine morphologies used for quantification with a 100x objective. Bottom, Representative pictures of CCI in CA1 dendrite and spine morphologies used for quantification with a 100x objective. ***B***, Top, Representative pictures of ACI in DG dendrite and spine morphologies used for quantification with a 100x objective. Bottom, Representative pictures of ACI in CA1 dendrite and spine morphologies used for quantification with a 100x objective. ***C***, Spine density in DG and CA1 after CCI. ***D***, Spine density in DG and CA1 after ACI. ***E***, Total dendritic length in DG and CA1 after CCI. ***F***, Total dendritic length in DG and CA1 after ACI. ***G***, Density of different spine types after CCI in DG. ***H***, Top, Examples of long thin, thin, stubby, and mushroom spines in DG after CCI. Bottom, Quantification classification of different morphological spine types. *p<0.05, **p<0.01, ***p<0.001 compared with the corresponding AAV-mCherry group.

Two-tailed *t*-test with Welch’s correction of spine densities identified significant differences in DG between the AAV-mCherry group and the AAV-hM4D(Gi)-mCherry group (**Fig. 5C**, CCI: t=3.757, p=0.0027; **Fig. 5D**, ACI: t=2.704, p=0.0192), suggesting a decrease in spine density with chemogenetic inhibition of DG hilar Sst^+^ interneurons. A significant decrease was observed in the AAV-hM4D(Gi)-mCherry group compared to the AAV-hM4D(Gi) group in the apical CA1 region only after CCI (t=3.698, p=0.0027), but not after ACI (t=1.345, p=0.1962) **(Fig. 5C,D)**. However, total dendritic length was significantly reduced after ACI and CCI (**Fig. 5 E**, DG in CCI: t=3.435, p=0.009; CA1 in CCI: t=2.065, p=0.0410; **Fig. 5 F**, DG in ACI: t=3.328, p=0.0014; CA1 in ACI: t=2.166, p=0.0327), indicating a reduction in total dendritic length in DG and apical CA1 after chronic and acute inhibition of DG hilar Sst^+^ interneurons.

Dendritic spines are often categorized by their various morphological subpopulations, such as stubby, mushroom, thin, and long thin (**Fig 5H**). We found a significant reduction of mushroom spines with other morphological types of spines unchanged (**Fig 5G**; two-way ANOVA, group × spine type, F_(3,40)_=2.25, Diff=-2.126, t=2.737, p<0.05). CCI resulted in a reduced number of mushroom spines, which are associated with long-term memory storage _29_.

### Cognitive function

Cognitive function has previously been reported to decline with aging in mice ^30^. To examine the impact of reduced activity of Sst^+^ interneurons on cognition, we performed the novel object recognition task (NOR), the Morris water maze test (MWM) including reversal learning, and the Y-Maze task (YM) in both ACI and CCI experiments **(Fig. 1B)**. It has been demonstrated before that performance in the NOR and the MWM is linked to CA1 ^31–34^, whereas reversal learning is linked to the dentate gyrus ^14^.

To investigate the potential spatial memory deficits in the experimental mice, we performed the MWM (CCI: n=11-15; ACI: n=15; **Fig. 6**). A two-way mixed ANOVA followed by Bonferroni post hoc test was conducted to investigate the impact of “genotype” (AAV-hM4D(Gi)-mCherry v.s. AAV-mCherry) and “day” on two behavioral measures, path length and latency to find the hidden platform during learning (L) and reversal learning phases (RL). In CCI (**Fig. 6A-F**), the AAV-hM4D(Gi)-mCherry mice showed significantly increased path length (<u>L:</u>day, day × genotype, F_(5,76)_=1.547, p < 0.01; t_(7)_=2.773, p < 0.05 for day 3. <u>RL:</u>day, day × genotype, F_(4,76)_=1.575, p < 0.01; t_(5)_=4.540, p < 0.001 for day 9; t_(5)_=2.837, p < 0.05 for day 11, t_(5)_=3.547, p < 0.01 for day 13) (**Fig. 6A**) and longer latency to find the hidden platform (<u>L:</u>day × genotype, F_(5,76)_=1.591, p < 0.01; t_(7)_=2.813, p < 0.05 for day 6; t_(7)_=4.70, p < 0.001 for day 7. RL:day × genotype, F_(4,76)_=2.634, p < 0.0001; t_(5)_ > 2, p < 0.01 for days 9,10,12,13) (**Fig. 6B**). In ACI, the AAV-hM4D(Gi)-mCherry mice did not differ from AAV-mCherry mice in their performance; no significant difference in latency nor path length to find the hidden platform were observed in AAV-mCherry and AAV-hM4D (Gi)-mCherry mice (path length: F_(1,94)_=0.8596, p > 0.05, **Fig. 6G**; latency to find the hidden platform: F_(1,94)_=1.082, p > 0.05, **Fig. 6H**).

**Figure 6.**
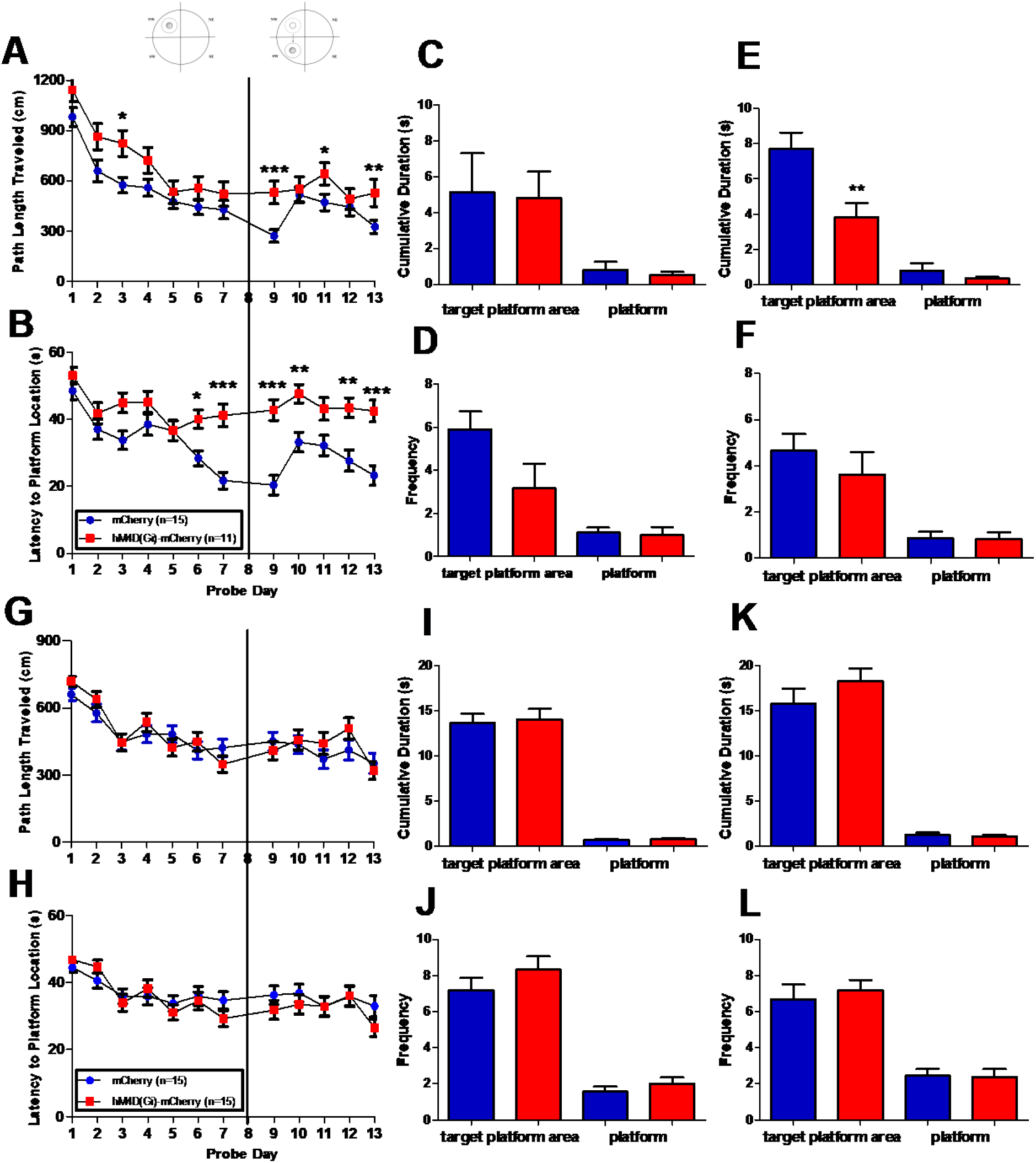
Morris water maze in AAV-mCherry and AAV-hM4D(Gi)-mCherry mice. Days1-7, learning phase; day8, learning phase probe trials; days9-13, reversal learning phase; day14, reversal learning probe trials. ***A-F***, Chronic Chemogenetic Inhibition (CCI). ***A***,***B***, Path length and latency to the hidden platform. ***C***,***D***, Cumulative duration and frequency to enter target platform area or the platform in learning phase probe trials. ***E***,***F***, Cumulative duration and frequency to enter target platform area or the platform in reversal learning phase probe trials. ***G-L***, Acute Chemogenetic Inhibition (ACI). ***G***,***H***, Path length and latency to hidden platform. ***I***,***J***, Cumulative duration and frequency to enter target platform area or the platform in learning phase probe trials. ***K***,***L***, Cumulative duration and frequency to enter target platform area or the platform in reversal learning phase probe trials. *p<0.05, **p<0.01, ***p<0.001.

After the initial learning phase and after the reversal learning phase, we conducted probe trials to determine memory retrieval ability in experimental mice. In the CCI learning probe trial, the AAV-hM4D(Gi)-mCherry mice displayed no significant difference for cumulative duration spent in the target platform area (unpaired t-test, p > 0.05, **Fig. 6C**) or frequency to enter the target platform area (unpaired t-test, p > 0.05, **Fig. 6D**). In the CCI reversal learning probe trial, the AAV-hM4D(Gi)-mCherry mice spent a significantly shorter time than the AAV-mCherry mice in the target quadrant (**Fig. 6E**) (unpaired t-test, AAV-hM4D(Gi)-mCherry: 7.708 ± 0.9183 n=11; AAV-mCherry: 3.825 ± 0.8063 n=15; F_(10,14)_ =1.769, p =0.3680). The frequency to enter the target quadrant was not different (unpaired t-test, p > 0.05, **Fig. 6F**).

In the ACI learning probe trial, the AAV-hM4D(Gi)-mCherry mice displayed no significant difference for cumulative duration in the target platform area and the frequency to enter the target platform area (unpaired t-test, p > 0.05, **Fig. 6I, J**). In the ACI reversal learning probe trial, the AAV-hM4D(Gi)-mCherry mice displayed no significant difference for cumulative duration and frequency to enter the target platform area (unpaired t-test, p > 0.05, **Fig. 6K**,**L**). The performance in the MWM learning and reversal learning phases revealed that the experimental mice had impaired performance only under CCI, while the ACI treated experimental mice showed an intact performance with no significant differences found in either path length or latency to find the hidden platform during learning, reversal learning, and probe trials.

We then determined whether loss of Sst^+^ interneurons in the DG hilus is associated with a decline in short-term recognition memory by analyzing AAV-hM4D (Gi)-mCherry and AAV-mCherry mice during both CCI and ACI in the NOR test. A significant decrease in the recognition index was observed in CCI (AAV-hM4D (Gi)-mCherry: 62.65 ± 3.406, n=15; AAV-mCherry: 48.12 ± 4.352, n=11; p=0.0125, t=2.670, df=28, unpaired t-test, **Fig. 7A**). In contrast, both ACI AAV-hM4D (Gi)-mCherry mice and ACI AAV-mCherry mice spent a similar amount of time in the proximity of the novel object (hM4D (Gi)-mCherry: 51.77 ± 3.489, n=13; mCherry: 59.63 ± 2.752, n=12; p > 0.05, unpaired t-test; **Fig. 7B**), suggesting a decreased recognition memory in CCI but not in ACI AAV-hM4D (Gi)-mCherry mice.

**Figure 7.**
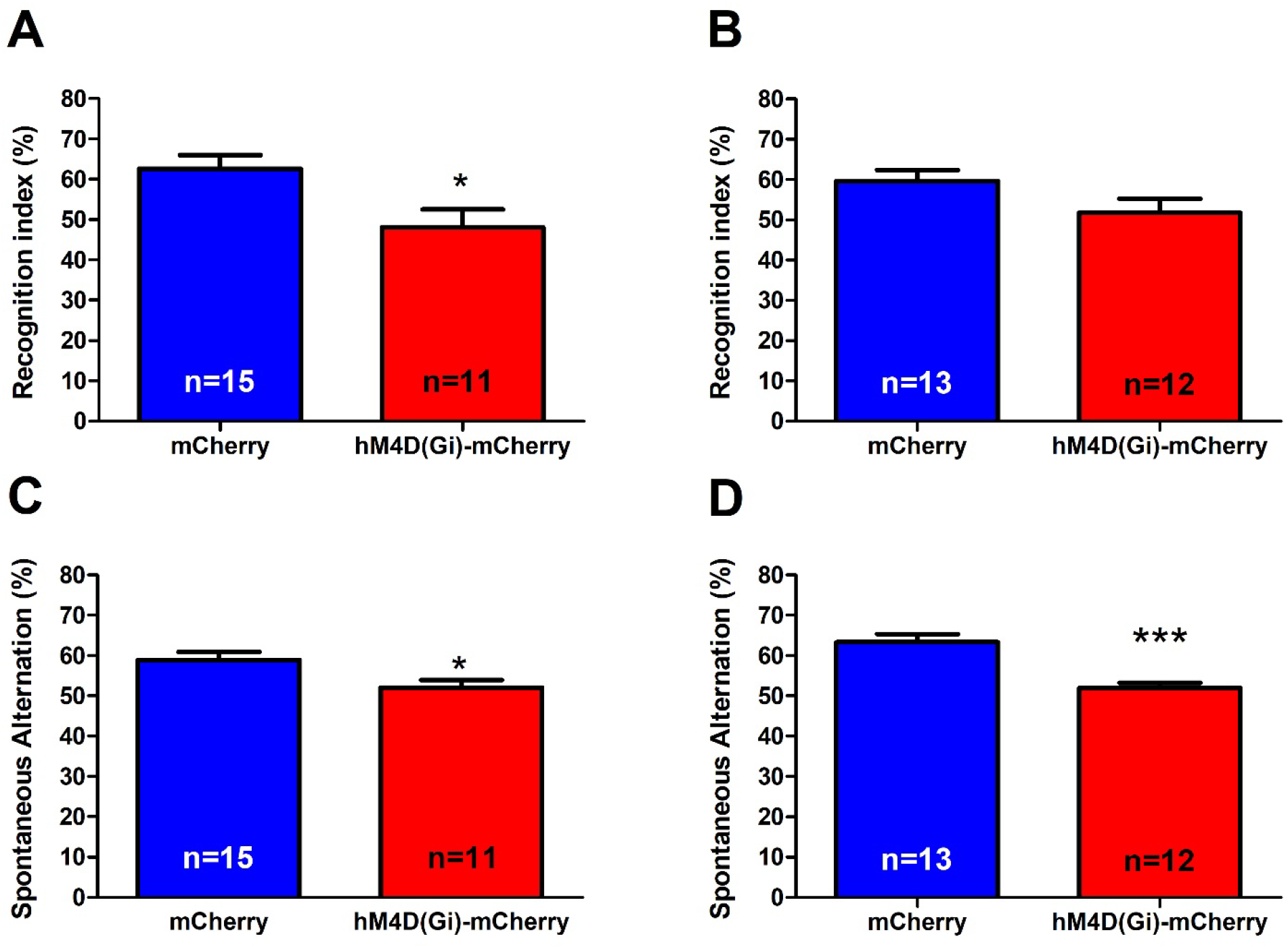
Novel Object Recognition Task (NOR) and Y-maze Task (YM) in AAV-mCherry and AAV-hM4D(Gi)-mCherry mice. ***A***, Recognition index in NOR in CCI. ***B***, Recognition index in NOR in ACI. ***C***, Spontaneous arm alternation percentage in YM in CCI. ***D***, Spontaneous arm alternation percentage in YM in ACI. *p<0.05, **p<0.01, ***p<0.001.

The Y-maze task was used to assess short-term working memory. In the CCI condition, AAV-hM4D(Gi)-mCherry mice had a significantly lower spontaneous alternation rate (AAV-mCherry: 58.91 ± 2.005, n=15; AAV-hM4D (Gi)-mCherry: 52.06 ± 1.811, n=11; p=0.0102, t=2.457, df=28, unpaired t-test) (**Fig. 7C**). In ACI, the spontaneous alternation rate also differed between groups with a lower recognition index in AAV-hM4D(Gi)-mCherry mice (AAV-mCherry: 63.31 ± 2.004, n=13; AAV-hM4D (Gi)-mCherry: 51.89 ± 1.362, n=12; p<0.0001, t=4.806, df=32; unpaired t-test) (**Fig. 7D**). Thus, the experimental (AAV-hM4D (Gi)-mCherry) mice had a significantly lower Y-maze spontaneous alternation rate than control (AAV-mCherry) mice in both ACI and CCI conditions.

## DISCUSSION

Understanding the cellular mechanisms underlying aging is crucial to develop novel strategies to prevent or to reverse age-related cognitive dysfunction. Based on previous studies in aged rats, which showed that expression of both Sst and GAD-67 proteins in DG hilus interneurons is reduced with age ^6^ and that the reduction of DG hilar Sst^+^ interneurons is correlated with cognitive impairments ^6,35^, we investigated two main questions: 1) whether a reduced activity of Sst^+^ interneurons in the DG hilus is sufficient to alter the activity of principal neurons in hippocampal subregions, to lead to changes in the expression of cellular markers and to impair of cognitive functions, and, if this is the case, 2) whether chronic (vs. acute) inhibition is required for these effects.

### Chronic chemogenetic inhibition of Sst^+^ hilar interneurons

Our experiments with chronic chemogenetic inhibition (CCI) of Sst^+^-positive interneurons in the DG hilus are summarized in **Table 1**. CCI resulted in an increase in the number of c-Fos-positive neurons in the DG, but not in CA3 or CA1 (**Fig. 2**), a reduction in the number of GAD67^+^ and Sst^+^ interneurons in the DG, but not in CA3 or CA1 (**Fig. 3**) and an increase in the IBA1^+^ cell density indicating microglial activation in DG, CA3, and CA1 (**Fig. 4**). The dendritic spine density was reduced in both DG and CA1 (**Fig. 5**). Behaviorally, learning and reversal learning in the MWM were impaired (**Fig. 6**), as were novel object recognition and alternation in the Y maze (**Fig. 7**). The results clearly demonstrate that chronic chemogenetic inhibition of Sst^+^-positive interneurons in the DG hilus is sufficient to induce cellular changes and cognitive dysfunction.

**Table 1.** Effects of chronic chemogenetic inhibition of DG hilus Sst^+^ interneurons (CCI) and acute chemogenetic inhibition of DG hilus Sst^+^ interneurons (ACI). The displayed results are comparing outcomes in AAV-hM4D(Gi)-mCherry mice compared to AAV-mCherry mice.

|  | **c-Fos<sup>+</sup>** | **Sst<sup>+</sup>** | **GAD-67<sup>+</sup>** | **Microglia**<br>**activation** | **MWM** | **NOR** | **YM** |
| --- | --- | --- | --- | --- | --- | --- | --- |
| **CCI** | ↑ | ↓ | ↓ | ↑ | ↓ | ↓ | ↓ |
| **ACI** | ↑ | — | — | — | — | — | ↓ |

### Acute chemogenetic inhibition of Sst^+^ hilar interneurons

When studying the effects of acute chemogenetic inhibtion (ACI) of Sst^+^-positive interneurons in the DG hilus (summarized in **Table 1**), we found an increase in the number of c-Fos-positive neurons in the DG, but not in CA3 or CA1, similar to what we observed with CCI (**Fig. 2**). In contrast to CCI, ACI did not reduce the number of GAD67^+^ and Sst^+^ neurons (**Fig. 3**), and, also in contract to CCI, ACI did not increase IBA1^+^ cell density in DG, CA3 and CA1, i.e., apparently did not induce microglial activation (**Fig. 4**). ACI lead to a reduction of dendritic spine density in DG (as with CCI) but not in CA1 (unlike CCI), and to a reduction of total dendritic length in both DG and CA1 (as with CCI) (**Fig. 5**). Behaviorally, unlike CCI, ACI had no effect on learning and reversal learning in the MWM (**Fig. 6**) and in the NOR test (**Fig. 6**), but like CCI, ACI reduced the spontaneous alternations in the YM (**Fig. 7**).

### Comparison of chronic and acute chemogenetic inhibition

Our studies revealed that while both ACI and CCI result in activation of neurons in the DG, a reduction in the number of GAD67^+^ and Sst+ interneurons in the dentate gyrus is only observed with CCI, not with ACI. Likewise, microglial activation in DG, CA3 and CA1 was only observed with CCI and not with ACI. Finally, spatial memory and novel object recognition memory were impaired only with CCI, but not with ACI. Only when testing working memory in the YM, both ACI and CCI reduced the percentage of correct alternations, and both ACI and CCI led to reductions in spine density and dendritic length. One potential interpretation for this is results is that while chronic inhibition is required to develop the full phenotype observed after CCI, the ACI protocol also has a “chronic” component, as clozapine is administered daily for 18 days. Notably, the experiments in which ACI induced some of the changes seen with CCI, the Y-Maze test is performed after 17 days of once daily treatment with clozapine and the cellular data after 18 days. In any case, the cellular data show some clear differences between ACI and CCI, indicating that these treatment protocols were substantially different. Moreover, the clear differences in some assays demonstrate that chronic inactivation of Sst^+^ interneurons is required for key elements of the observed phenotype with CCI. Overall, these results strongly support the hypothesis that chronic inhibition of Sst^+^ interneurons in the DG hilus is necessary and sufficient for microglial activation and cognitive dysfunction. This dependence on chronic inhibition of these neurons for inducing cognitive dysfunction also aligns with the chronicity of the aging process, supporting the view that a gradual decline of function of hilar Sst^+^ interneurons, as likely occurs with age-related reduction in the number of hilar Sst+ interneurons ^6,35^, contributes to age-related cognitive decline. Indeed, when considering chronic chemogenetic inhibition as a potential model for hippocampal aging, our results are in line with previous studies which showed memory impairments in MWM, YM and NOR in aged rodents ^36–38^, indicating that memory impairment and aging are closely linked and may be dependent on the same hippocampal neural circuits ^14,32,39^.

While the current study was underway, Fee et al. (2021) reported a study on the role of Sst^+^ cells in regulating mood and cognitive functions ^40^. They used a global approach and developed a chemogenetic model which inhibits Sst^+^ interneuron function brain-wide by injecting their chemogenetic inhibitory receptor-containing viral construct intraventricularly. They employed clozapine-N-oxide (CNO) to activate the DREADD instead of clozapine used in the current study and also performed the YM and NOR tasks. They achieved repeated acute silencing of Sst^+^ cells by administrating CNO 30 minutes before each behavioral experiment, which is similar to our ACI model. While they found that their CNO-treated mice spent significantly less time with the novel object than with the familiar object in the NOR task and there were no differences between groups in the YM task, with acute injections of clozapine we found no difference between AAV-mCherry-injected mice and AAV-hM4D(Gi)-mCherry-injected mice in the NOR task, but AAV-hM4D(Gi)-mCherry-injected mice displayed significantly fewer spontaneous alternations in the YM task. Although the reasons for the observed differences are not understood, there are some potential explanations, including differences in dosing (3.5 mg/kg CNO per day in Fee et al., 2021 vs. 0.1 mg/kg clozapine per day in our ACI and CCI studies) ^40^, multiple experiments being performed with repeated acute dosing which might result in chronic-like effects especially at higher drug doses, and their generalized Sst^+^ interneuron inhibition could affect multiple neural circuits while we inhibited a narrowly targeted cell population. In any case, both studies demonstrated effects of Sst^+^ cell regulation on cognitive functions. Interestingly, a difference between acute and chronic chemogenetic silencing of Sst^+^ interneurons has also been reported by Soumier et al. ^41^, who described that while acute chemogenetic inhibition of Sst^+^ interneurons in the frontal cortex resulted in increased behavioral emotionality, chronic chemogenetic inhibition resulted in decreased behavioral emotionality.

Several lines of evidence point to a role of GABAergic inhibition for hippocampal learning and memory. In mice in which hilar GABAergic interneurons were silenced optogenetically, c-Fos^+^ expression in the DG (but not in CA3 or CA1) and the firing rate of the GCs were increased, resulting in impaired spatial learning and memory retrieval ^7^. Aged rats with memory impairment had restoration of hilar Sst expression after receiving levetiracetam, a compound that modulates synaptic neurotransmitter release ^6^. Mice with decreased GABA levels in the hippocampus induced by overexpressing GABA transporter 1, which mediates GABA reuptake, display deficits in learning and memory ^42^. A lack of α5-GABA_A_Rs in the DG granule cells, which presumably led to a reduced inhibitory input into the granule cells, has previously been shown to result in a reduced tonic inhibition of DG granule cells and reversal learning deficits in the MWM ^14^. All of these studies thus revealed a critical role of GABAergic neurotransmission for hippocampal learning and memory. Thus, our current study demonstrating that inhibition and thus reduced activity of a GABAergic interneuronal subtype results in cognitive deficits is line with previous evidence demonstrating essential roles for GABA in learning and memory. Moreover, GABA levels in the mouse brain as determined by ^1^[H] magnetic resonance spectroscopy are reduced with aging ^43^. Moreover, increased microglial activation is known to be associated with human aging ^44^. In our study, we found microglial activation in DG, CA3, and CA1 regions in CCI mice, indicating the inactivation of Sst^+^ interneurons causes changes in the hippocampus that eventually result in an increase of this microglial activation marker.

### Conclusion and Future Directions

In summary, our experiments demonstrate a causal relationship between a chronic loss of activity of dentate hilus Sst^+^ interneurons, microglial activation, and cognitive dysfunction. Given that a reduction in the number of these interneurons has been found to be associated with age-related memory deficits ^6,35^, a reduced number or function of hilar Sst^+^ interneurons is thus likely to be an important factor in the development of age-related cognitive dysfunction. Unlike the acute chemogenetic inhibition model which also results in increased c-Fos staining and hyperactivity in DG, the chronic chemogenetic inhibition model mimics several relevant features of hippocampal aging and may thus be further evaluated as an experimental model of aging-related processes in the hippocampus, which also allows to study the potential reversibility of the phenotype after stopping clozapine administration. This novel model may also be useful to study postoperative neurocognitive disorder and other age-related cognitive disorders, and for the development of novel therapeutic approaches for such disorders.

## MATERIALS AND METHODS

### Animals

Adult Sst-IRES-Cre transgenic mice (Stock no. 013044, The Jackson Laboratory) were crossed with C57BL/6J mice (Stock no. 000664, The Jackson Laboratory). Hemizygous adult mice of both sexes (3-4 months) were used for all studies. All mice were housed in a climate-controlled room maintained on a 12:12 light-dark cycle (lights on at 7 a.m., lights off at 7 p.m.) with food and water ad libitum. All procedures are approved by the Institutional Animal Care and Use Committee at University of Illinois at Urbana-Champaign. ARRIVE guidelines were followed.

### Silencing Sst^+^ Cell Function in the DG Hilus

To achieve a Sst^+^ cell-specific manipulation, we used somatostatin-IRES-Cre (Sst-Cre) mice, which express Cre recombinase in somatostatin-expressing neurons. We bred heterozygous Sst-Cre mice with wildtype mice to generate experimental Sst-Cre mice. A G_i_-coupled <u>d</u>esigner <u>r</u>eceptor <u>e</u>xclusively <u>a</u>ctivated by <u>d</u>esigner <u>d</u>rugs (DREADD) was used that decreases the overall amount of cAMP, which results in neural inhibition ^21,22^. To achieve region-selective manipulation, we stereotaxically injected a viral G_i_-DREADD construct that will only be expressed in Cre-positive cells into the dentate gyrus hilus region, where it can be activated with clozapine N-oxide (CNO) or clozapine. While CNO is likely the most widely used ligand for chemogenetic studies, clozapine displays higher affinity and greater potency for hM4D(Gi) ^22^. Clozapine has favorable pharmacokinetic properties for chronic administration ^23,24^ and was therefore used in our experiments.

### Surgery

In order to study the role of Sst^+^ in the dentate hilus, we generated mice in which the Sst^+^ in the dentate hilus can be silenced chemogenetically. Sst-Cre mice (3-4 months of age) received ketoprofen 5mg/kg, s.c. and atropine 0.04 mg/kg, s.c. before surgery. Mice were anesthetized with isoflurane (2-3%) and maintained under anesthesia (1.5%) throughout the surgery. AAV-hSyn-DIO-hM4D(Gi)-mCherry (Addgene #44362) was injected bilaterally into the DG hilus (stereotaxic coordinates AP: 2.1mm, ML: ±1.5mm, DV: -2.1mm relative to bregma) at a rate of 0.120 µL/min (1000nL per side), and the needle was left in place for an additional 2 mins to permit diffusion. The control groups were injected with an AAV vector, AAV-hSyn-DIO-mCherry (Addgene #50459), lacking the chemogenetic receptor hM4D(Gi). Injection locations were verified histologically at the end of the study.

Clozapine dihydrochloride (water soluble, Hello Bio #HB6129) was administered to the mice to activate hM4D(Gi) and thus inhibit hilar Sst^+^ interneurons. The chronic treatment and control groups received clozapine (0.1mg/kg/day) in the drinking water for 21 days before and throughout all testing and the acute treatment groups and control groups were administered clozapine (0.1mg/kg i.p.) 30 min before behavioral experiments started. Mice drink approximately 4 mL of water per day. Males weigh more than females (e.g., 35 g versus 25 g, so the males may get 3.5 ug/4ml and the females 2.5 ug/4ml). Mice were weighed every 2 days, and the clozapine solution adjustments made every 2 days to reach the desired concentration.

### Behavioral Tests

All behavioral experiments were performed during the light phase of the light/dark cycle. All mouse behaviors were monitored using the EthoVision XT video tracking system.

#### Novel Object Recognition Task

Mice were habituated in the experimental chamber for two days before the testing phase and each habituation period lasted for 15 min for each animal per session. Each mouse was presented with two identical objects for 10 min on the actual test day. After one hour, the mice were brought back to the same chamber and presented with one of the same training objects and one novel object. Interaction time was recorded using the multiple body point module. Mice were considered exploring the novel object when the nose point as defined by the EthoVision XT software was in close proximity to the object. A novelty recognition index was calculated by dividing the time spent in the proximity of the novel object by the total time spent with both objects.

#### Morris Water Maze and Reversal Learning

A round pool (diameter: 120 cm) is filled with a mixture of water (22-24°C) and a white, non-toxic dye (Blick Premium Grade Tempera). A 10cm-diameter platform was submerged 2cm under the water surface. A 25cm-diameter target platform area was defined in the same quadrant as a concentric circle around the target platform. The frequency to enter and cumulative duration spent in the target platform are also recorded and analyzed. Visual cues were in the four quadrants of the pool in shapes with different geometry. Mice performed three trials daily from day 1 to day 7, released from a different quadrant each time in random order while the target platform location was constant. A trial ended when the animal found and stayed on the platform for 2 seconds. When an animal failed to reach the platform within 60 seconds, the experimenter guided it to the platform and put it back to the cage after staying on the platform for 10 seconds. From day 10 to day 13, the reversal learning phase was established by moving the platform from the original location to the nearest quadrant to increase the effects of interference. During probe trial (day 8) and reversal learning probe trial (day 14), the platform was removed, and the mice were left in the pool for 120 seconds.

#### Y-Maze Task

The apparatus was a Y-shaped maze with three gray, opaque plastic arms at a 120° angle from each other. The arm length is 31cm and the arm width is 7cm. Mice were placed in the center of the Y-Maze and allowed to explore any of the three closed arms freely for 8 min. The spontaneous arm alternation rate is calculated by dividing the number of spontaneous alternations by the number of total arm entries and multiplication by 100. Mice with intact working memory would run in the three arms with spontaneous alternations, meaning that at each left-right decision point, they make an alternating decision. In the YM, a spontaneous alternation rate of 50% would imply random choices.

### Histology and Immunohistochemistry

Mice were deeply anesthetized with ketamine (139 mg/kg i.p.) and xylazine (21 mg/kg i.p.) and then transcardially perfused with ice-cold phosphate-buffered saline (PBS) followed by 4% ice-cold paraformaldehyde. After 24 hours of post-fixation at 4°C, brains were moved into 30% sucrose for 72 hours and sectioned into 30 µm coronal sections using a cryostat. Free-floating sections were collected mounted on slides for staining.

We performed analysis of sections for target cell labeling and quantification. For analysis of c-Fos staining, all mice (n=6 per group) were exposed to a novel environment for 10 min and then placed into a clean cage individually for 1 hour before perfusion. For other stainings, free-floating sections (n=3 per sex, n=6 per group) were washed in 0.1 M phosphate-buffered saline (PBS) to remove cryoprotectant. The sections were blocked in blocking solution (2% NGS and 0.4% Triton X-100 in PBS) for 2 hours at room temperature, then incubated in the primary antibody at 4°C overnight. The primary antibodies used were as follows: rabbit anti-c-Fos (Cell Signaling Tech, cat # 2250, 1:500), anti-somatostatin (SOM, Invitrogen, cat # PA5-82678, 1:1000), anti-glutamic acid decarboxylase 67 (GAD-67, Invitrogen, cat # PA5-21397, 1:500), anti-IBA1 on microglia (IBA1, Invitrogen, cat # MA5-36257, 1:500), anti-neuronal nuclei (NeuN, Cell Signaling Tech, cat # 24307, 1:400). After washing in blocking solution and 3% H_2_O_2_, the sections were incubated in a biotinylated goat anti-rabbit secondary antibody in blocking solution (Invitrogen, cat # 31820, 1:500) and transferred in a detection reagent (Vecstatin Elite ABC kit, Vector Laboratories, cat # PK-7200). The sections were incubated in a 3,3’-diaminobenzidine (DAB) solution. After rinsing, sections were mounted, dehydrated, and cover-slipped with mounting medium (Eukitt Quick-handling). The section slides were imaged with an Olympus BX51 microscope. To verify viral placements and immunofluorescence labeling of mCherry, DAPI stain (Abcam, cat# ab228549) was added to free-floating 30 µm coronal sections using the concentration of 1µM in the dark. The sections were mounted and cover-slipped on slides using Eukitt Quick-handling mounting medium, and imaged with a Leica DM2500 microscope.

### Golgi-Cox Staining

Mouse brains were removed from skulls without fixation. The fresh brain tissue was immersed in the impregnation solution (superGolgi Kit, Bioenno Tech) for 12 days and stored at 4°C in the dark. After being transferred to and incubated in the post-impregnation buffer for 2 days, the brain tissues were sectioned into 150µm thick sections using a vibratome. The sections were collected and mounted on gelatin-coated slides, dehydrated, and cleaned. Then they were cover-slipped using a mounting medium (Sigma-Aldrich) and stored at room temperature in the dark.

### Dendritic Spine Analysis

We examined spines on apical dendrites of CA1 pyramidal neurons and dendrites of the dorsal dentate gyrus. To ensure the accurate measurements, we only evaluated dendrites that showed no breaks in the staining ^25^ and that were not interrupted by other neurons or artifacts ^26^. Primary spines were not analyzed, we only evaluated spines located on secondary or tertiary dendritic trees. One segment per individual dendritic branch and two branches per neuron were chosen for the analysis. Quantitative two-dimensional analyses of dendritic and spine fragments were conducted by using a stereological microscope (Zeiss AxioImager A1 light microscope) with a 100x objective (oil immersion). For each dendrite at least 3 images along each spine segment were taken and spine densities calculated. Spine densities per 10µm and spine length from the dendrite shaft to the spine head were marked and calculated by using *Reconstruct* software (Version 1.1.0.1) ^27^.

### Statistical Analysis

Statistical comparisons were performed using Graph Pad Prism (Graph Pad Software Inc., La Jolla, CA). In the c-Fos, SOM, GAD-67, and NeuN staining experiments, the average numbers of c-Fos^+^ nuclei, Sst^+^, GAD-67^+^ cells, and NeuN^+^ cells in the DG, CA3, and CA1 regions were analyzed using two-way repeated-measures analysis of variance (ANOVA). Each slide was counted three times, and the mean value was used in the ANOVAs. IBA1 staining was used as a marker for microglial activation. The average IBA1^+^ density was analyzed with one-way ANOVA followed by Newman-Keuls multiple comparison tests. Statistical differences between treatments and regions were assessed using Bonferroni’s post-hoc comparison test. Performance in behavioral experiments was be analyzed using one-way ANOVA and Scheffe’s test.

## ACKNOWLEDGEMENTS

Research reported in this paper was supported by the National Institute of General Medical Sciences of the National Institutes of Health under award number R01GM128183 to U.R.. The content is solely the responsibility of the authors and does not necessarily represent the official views of the National Institutes of Health.

## DATA AVAILABILITY

All data are included in the manuscript and/or Supplementary Material.

## Notes

### Competing Interest Statement

UR serves on the Scientific Advisory Board of Damona Pharmaceuticals.

### Summary of Updates

The language of the text has been updated and the title has been changed.

